# Ultraviolet radiation exposure and melanoma: evidence for gene-environment interaction in a large prospective cohort

**DOI:** 10.1101/666123

**Authors:** Catherine M. Olsen, Nirmala Pandeya, Matthew H. Law, Stuart MacGregor, Mark M. Iles, Bridie S. Thompson, Adele C. Green, Rachel E. Neale, David C. Whiteman, for the QSkin Study

**Affiliations:** Department of Population Health, QIMR Berghofer Medical Research Institute, Queensland, Australia; Faculty of Medicine, University of Queensland, Queensland, Australia; School of Public Health, University of Queensland, Queensland, Australia; Statistical Genetics, QIMR Berghofer Medical Research Institute, Queensland, Australia; Leeds Institute of Medical Research at St James’s, University of Leeds, Leeds, UK & Leeds Institute for Data Analytics, University of Leeds, Leeds, UK; Cancer Research UK Manchester Institute and University of Manchester, Manchester Academic Health Science Centre, Manchester, UK

## Abstract

Melanoma develops as the result of complex interactions between sun exposure and genetic factors. Data on the relationship between sunlight and melanoma from prospective studies are scant, and the combination of ultraviolet exposure data collected before melanoma diagnosis and genetic information is rarer still. We aimed to quantify the association between ambient and personal UV exposure in relation to risk of incident melanoma (invasive; invasive+*in situ*) in a large population-based prospective study of men and women (n=38,833) residing in a high ambient UV setting, and to examine potential gene-environment interactions. During a median follow-up time of 4.4 years, 782 (1.5%) participants developed cutaneous melanoma (316 invasive, 466 *in situ*). Country of birth, age at migration and sunburns during all periods of life were significantly associated with melanoma risk. Histories of keratinocyte cancer and of other actinic lesions were both strongly associated with melanoma risk. An interaction with polygenic risk is possible; among people at low risk, markers of cumulative sun exposure were associated with melanoma. In contrast, among people at high polygenic risk, markers of high-level early life ambient exposure were associated with melanoma. Polygenic risk scores can assist in identifying individuals for whom sunlight exposure is most relevant.

## INTRODUCTION

Ultraviolet (UV) radiation is the only environmental factor that has been consistently implicated as a cause of melanoma, and is estimated to account for between 63 and 90% of melanoma cases (Armstrong and Kricker, 1993; Olsen *et al.*, 2015). The relationship is complex, however, and exposure effects are highly modified by host factors and behaviors. Associations with an intermittent pattern of sun exposure and a history of sunburns have been reported consistently, but not with occupational exposure or a high continuous pattern of sun exposure (Gandini *et al.*, 2005). However, much of the extant literature on the association derives from case-control studies, which have inherent limitations including selection and recall bias. Of the 57 studies included in the most recent systematic review (published in 2005), five were cohort studies; however, these were all occupational cohort studies, where occupation was used as a proxy for sun exposure measures (i.e. indoor/outdoor work), and all were retrospective in nature (Gandini *et al.*, 2005). Cohort data published subsequent to the systematic review are limited to four studies in women (Cho *et al.*, 2005; Han *et al.*, 2006; Nielsen *et al.*, 2012; Wu *et al.*, 2016; Ghiasvand *et al.*, 2018; Savoye *et al.*, 2018) and one in men (Cho *et al.*, 2005; Wu *et al.*, 2016); for one of these, information on sun exposure was only collected for a subset of the cohort (Savoye *et al.*, 2018), and for others, information on important potential confounders was not collected at baseline (Cho *et al.*, 2005; Han *et al.*, 2006; Wu *et al.*, 2016). Significant associations were reported with various definitions of sunburns (Cho *et al.*, 2005; Han *et al.*, 2006; Nielsen *et al.*, 2012; Wu *et al.*, 2016) and ‘sunny vacations’ (Cho *et al.*, 2005), but not with measures of cumulative sun exposure (Savoye *et al.*, 2018) or region of residence (Han *et al.*, 2006; Wu *et al.*, 2016).

Quantifying individual exposure to UV radiation in observational studies is difficult. Measures of sun exposure vary markedly across studies, and generally have low reproducibility (English *et al.*, 1998; van der Mei *et al.*, 2006). Markers of actinic damage, such as actinic keratoses and keratinocyte cancers, can act as proxies of high cumulative UV dose received at the dermoepidermal junction. The relationship between personal history of keratinocyte cancer and risk of melanoma has been sparsely studied using prospective data (Wu *et al.*, 2017), and it is hypothesized that the relationship may differ for melanomas arising on different body sites (Whiteman *et al.*, 2003).

The role of genotype on the relationship between UV radiation exposure and melanoma risk is also poorly understood. An examination of the potential interaction between genetic factors and UV radiation exposure might elucidate the biological processes that lead to melanoma, and may help identify those people for whom UV exposure is most hazardous, especially those with no high-risk phenotypic features.

We aimed to address these issues by examining the association between measures of UV radiation exposure (including measures of actinic damage) and incident melanoma (invasive only; and invasive + *in situ*) in a large population-based prospective study of men and women residing in a region of high insolation, including an examination of potential mediating effects and gene-environment interactions.

## RESULTS

Of 38,833 eligible participants, 21,067 (54%) were women and the mean age was 56 years (SD 8.2). During a median (and mean) follow-up time of 4.4 years, 783 (1.5%) participants developed melanoma (317 invasive melanomas, 507 *in situ* melanomas); 41 developed both invasive and *in situ* melanoma. Of the 317 incident invasive cases, 14 were diagnosed with *in situ* melanoma after enrolment but prior to their invasive melanoma diagnosis. Melanoma cases were more likely to be male (59% vs 41%; p<0.001) and on average, were older than non-cases at baseline (58.6 years vs. 55.9 years, respectively; p<0.001). Of the invasive cases, 61% were of the superficial spreading subtype and 76% were <1mm (Table 1). One case of acral melanoma was excluded, leaving 316 invasive cases.

**Table 1.** Tumor characteristics of first primary melanoma diagnosed during follow-up (n=782).

| Characteristic | Invasive (n=316) | *In situ* (n=466) |
| --- | --- | --- |
|  | N (%) | N (%) |
| **Age** |  |  |
| 40-49 | 46 (14.6) | 83 (17.8) |
| 50-59 | 116 (36.7) | 160 (34.3) |
| ≥60 | 154 (48.7) | 223 (47.9) |
| **Sex** |  |  |
| Female | 136 (43.0) | 185 (39.7) |
| Male | 180 (57.0) | 231 (60.3) |
| **Body site** |  |  |
| Head/neck | 48 (15.2) | 115 (24.7) |
| Trunk | 110 (34.8) | 166 (35.6) |
| Upper limbs | 83 (26.3) | 114 (24.5) |
| Lower limbs | 64 (20.3) | 57 (12.2) |
| Missing | 11 (3.5) | 14 (3.0) |
| **Thickness category** |  |  |
| <1 mm | 240 (76.0) | n.a. |
| 1-1.99 mm | 39 (12.3) |  |
| 2-3.99 mm | 18 (5.7) |  |
| ≥4 mm | 9 (2.8) |  |
| Missing | 10 (3.2) |  |
| **Clarks level** |  |  |
| 1 |  | 466 (100) |
| 2 | 165 (52.2) |  |
| 3 | 68 (21.5) |  |
| 4 | 64 (20.3) |  |
| 5 | 6 (1.9) |  |
| Missing | 13 (4.0) |  |
| **Melanoma type** |  |  |
| Superficial spreading | 194 (61.4) | 141 (30.3) |
| Lentigo maligna | 17 (5.4) | 150 (32.2) |
| Nodular | 22 (7.0) |  |
| Other<sup>1</sup> | 83 (26.3) | 175 (37.6) |

<sup>1</sup>Other for invasive melanoma includes: amelanotic melanoma, malignant melanoma in junctional nevus, desmoplastic malignant melanoma, regressing malignant melanoma, spindle cell melanoma not otherwise specified and malignant melanoma not otherwise specified. Other for *in situ* melanoma includes: melanoma in situ and junctional nevus in situ.

### Invasive melanoma

People born in Australia had a significantly higher risk than those born elsewhere, as did those born at latitudes <45° N/S (Table 2). There was no significant association between the region of Australia in which participants lived longest as a child or over the entire lifetime. Compared to native-born participants, those born overseas who moved to Australia when aged 20 years or older had a significantly lower risk (OR 0.48, 95% CI 0.29-0.78).

**Table 2.** Measures of ambient sun exposure and risk of first invasive melanoma.

| Variables | Participants with invasive melanoma |  | Crude<br>HR (95%CI) | Age & sex adj<br>HR (95%CI) |
| --- | --- | --- | --- | --- |
|  | No (n=38,516) | Yes (n=316) |  |  |
|  | n (%) | n (%) |  |  |
| **Born in Australia** |  |  |  |  |
| No | 6782 (17.6) | 41 (13.0) | Reference | Reference |
| Yes | 31,734 (82.4) | 275 (87.0) | 1.43 (1.03-1.98) | 1.53 (1.10-2.13) |
| **Latitude of birth** |  |  |  |  |
| ≥45° S/N | 4503 (11.7) | 26 (8.2) | Reference | Reference |
| <45° S/N | 34,013 (88.3) | 290 (91.8) | 1.47 (0.99-2.20) | 1.62 (1.09-2.42) |
| **Region of Australia lived longest up to age 20 years** |  |  |  |  |
| Southern region | 11,646 (33.8) | 101 (34.1) | Reference | Reference |
| Central region | 17,033 (49.4) | 159 (53.7) | 1.09 (0.85-1.41) | 1.13 (0.88-1.46) |
| Northern region | 5814 (16.9) | 36 (12.2) | 0.73 (0.50-1.06) | 0.76 (0.52-1.12) |
| **Region of Australia lived longest over lifetime** |  |  |  |  |
| Southern region | 8776 (22.8) | 74 (23.4) | Reference | Reference |
| Central region | 22,899 (59.5) | 202 (63.9) | 1.03 (0.79-1.35) | 1.06 (0.81-1.38) |
| Northern region | 6841 (17.8) | 40 (12.7) | 0.69 (0.42-1.02) | 0.70 (0.48-1.04) |
| **Age moved to Australia** |  |  |  |  |
| Born in Australia | 31,774 (82.5) | 275 (87.0) | Reference | Reference |
| 1-10 years | 1874 (4.9) | 12 (3.8) | 0.74 (0.41-1.32) | 0.73 (0.41-1.29) |
| 11-20 years | 1092 (2.8) | 12 (3.8) | 1.27 (0.71-2.27) | 1.19 (0.66-2.12) |
| 20+ years | 3776 (9.8) | 17 (5.4) | 0.52 (0.32-0.85) | 0.48 (0.29-0.78) |

A history of greater than 50 sunburns as a youth and 21-50 sunburns as an adult was associated with an over two-fold increased risk (Table 3). Sunburns as a child were not significantly associated with melanoma risk after adjusting for hair color and skin tanning ability. Our mediation analyses suggest that 8.7% of the effect of sunburns in youth on melanoma risk is mediated via the influence of nevus density, and the mediation effect was significant (p<0.001). Although we found no evidence that melanoma was associated with self-reported cumulative sun exposure, average hours in the sun on weekdays or weekends, or sunbed use, we found significant 2-3 fold elevations in risk with the number of previous keratinocyte cancers and number of treatments for actinic lesions (Table 4).

**Table 3.** Measures of personal sun exposure and risk of first invasive melanoma.

| Variables | Invasive melanoma |  | Crude<br>HR (95%CI) | Age & sex adj<br>HR (95%CI) | Model 1<sup>1</sup><br>HR (95%CI) |
| --- | --- | --- | --- | --- | --- |
|  | No (n=38,516)<br>n (%) | Yes (n=316)<br>n (%) |  |  |  |
| Sunburns as a child (less than 10 years) |  |  |  |  |  |
| Never | 8081 (21.0) | 58 (18.4) | Reference | Reference | Reference |
| 1-5 times | 17,582 (45.7) | 149 (47.2) | 1.15 (0.84-1.57) | 1.17 (0.85-1.60) | 1.06 (0.77-1.45) |
| 6-10 times | 6732 (17.5) | 54 (17.1) | 1.05 (0.71-1.54) | 1.08 (0.73-1.60) | 0.93 (0.63-1.39) |
| 11-20 times | 3638 (9.5) | 27 (8.5) | 1.08 (0.69-1.70) | 1.15 (0.73-1.83) | 0.94 (0.59-1.51) |
| 21-50 times | 1748 (4.5) | 15 (4.8) | 1.22 (0.69-2.15) | 1.27 (0.72-2.24) | 1.01 (0.57-1.82) |
| >50 times | 735 (1.9) | 13 (4.1) | 2.24 (1.20-4.17) | 2.36 (1.26-4.41) | 1.81 (0.95-3.44) |
| Sunburns as a youth (10-20 years old) |  |  |  |  |  |
| Never | 1788 (4.6) | 9 (2.9) | Reference | Reference | Reference |
| 1-5 times | 16,794 (43.6) | 132 (41.8) | 1.55 (0.77-3.12) | 1.77 (0.88-3.58) | 1.59 (0.78-3.21) |
| 6-10 times | 9579 (24.9) | 83 (26.3) | 1.71 (0.84-3.48) | 2.03 (0.99-4.14) | 1.76 (0.86-3.60) |
| 11-20 times | 6416 (16.7) | 53 (16.8) | 1.59 (0.76-3.29) | 1.91 (0.92-3.97) | 1.60 (0.77-3.35) |
| 21-50 times | 2886 (7.5) | 21 (6.7) | 1.47 (0.66-3.28) | 1.78 (0.79-3.97) | 1.47 (0.65-3.30) |
| >50 times | 1053 (2.7) | 18 (5.7) | 3.17 (1.38-7.25) | 3.77 (1.64-8.65) | 3.04 (1.31-7.06) |
| Sunburns as an adult (more than 20 years old) |  |  |  |  |  |
| Never | 6230 (16.2) | 41 (13.0) | Reference | Reference | Reference |
| 1-5 times | 20,461 (53.1) | 164 (51.9) | 1.25 (0.88-1.77) | 1.34 (0.94-1.91) | 1.37 (0.96-1.95) |
| 6-10 times | 6396 (16.6) | 57 (18.0) | 1.33 (0.88-2.01) | 1.43 (0.94-2.18) | 1.47 (0.97-2.23) |
| 11-20 times | 3409 (8.9) | 26 (8.2) | 1.19 (0.72-1.96) | 1.25 (0.76-2.07) | 1.27 (0.77-2.10) |
| 21-50 times | 1462 (3.8) | 22 (7.0) | 2.32 (1.37-3.92) | 2.37 (1.40-4.02) | 2.35 (1.39-4.00) |
| >50 times | 558 (1.5) | 6 (1.9) | 1.65 (0.70-3.91) | 1.59 (0.67-3.77) | 1.63 (0.68-3.87) |
| Cumulative sun exposure<sup>2</sup> |  |  |  |  |  |
| Q1 | 9341 (24.3) | 72 (22.8) | Reference | Reference | Reference |
| Q2 | 9674 (25.1) | 67 (21.2) | 0.89 (0.63-1.25) | 0.80 (0.57-1.12) | 0.84 (0.60-1.18) |
| Q3 | 9728 (25.3) | 85 (26.9) | 1.09 (0.79-1.51) | 0.87 (0.62-1.21) | 0.94 (0.68-1.31) |
| Q4 | 9773 (25.4) | 92 (29.1) | 1.21 (0.88-1.66) | 0.78 (0.56-1.08) | 0.87 (0.62-1.21) |
| Average number of hours in the sun on weekdays<sup>2</sup> |  |  |  |  |  |
| Q1 | 9032 (23.5) | 72 (22.8) | Reference | Reference | Reference |
| Q2 | 9443 (24.5) | 76 (24.1) | 1.00 (0.72-1.39) | 0.98 (0.70-1.36) | 1.02 (0.73-1.41) |
| Q3 | 10,154 (26.4) | 82 (26.0) | 0.98 (0.71-1.37) | 0.88 (0.64-1.22) | 0.94 (0.68-1.30) |
| Q4 | 9887 (25.7) | 86 (27.2) | 1.08 (0.78-1.48) | 0.84 (0.60-1.16) | 0.91 (0.66-1.27) |
| Average number of hours in the sun on weekend days<sup>2</sup> |  |  |  |  |  |
| Q1 | 9694 (25.2) | 77 (24.4) | Reference | Reference | Reference |
| Q2 | 9857 (25.6) | 84 (26.6) | 1.03 (0.74-1.42) | 1.01 (0.73-1.40) | 1.08 (0.78-1.50) |
| Q3 | 9775 (25.4) | 74 (23.4) | 0.96 (0.69-1.33) | 0.88 (0.63-1.22) | 0.97 (0.70-1.35) |
| Q4 | 9190 (23.9) | 81 (25.6) | 1.07 (0.78-1.48) | 0.93 (0.66-1.30) | 1.05 (0.75-1.48) |
| Sunbed use |  |  |  |  |  |
| Never | 34,401 (89.3) | 285 (90.2) | Reference | Reference | Reference |
| Ever | 4115 (10.7) | 31 (9.8) | 0.91 (0.63-1.32) | 1.20 (0.82-1.74) | 1.31 (0.90-1.91) |

<sup>1</sup>Model 1- additionally adjusted for hair colour, tanning ability<sup>2</sup>Q - Quartile

**Table 4.** Proxy measures of personal sun exposure and risk of first invasive melanoma.

| Variables | Invasive melanoma |  | Crude<br>HR (95%CI) | Age & sex adj<br>HR (95%CI) | Model 1<sup>1</sup><br>HR (95%CI) |
| --- | --- | --- | --- | --- | --- |
|  | No (n=38,516)<br>n (%) | Yes (n=316)<br>n (%) |  |  |  |
| **Number of skin cancers excised** |  |  |  |  |  |
| None | 23,536 (61.1) | 122 (38.6) | Reference | Reference | Reference |
| 1 | 5367 (13.9) | 46 (14.6) | 1.63 (1.16-2.31) | 1.54 (1.09-2.17) | 1.46 (1.03-2.05) |
| 2-10 | 8050 (20.9) | 111 (35.1) | 2.67 (2.06-3.45) | 2.36 (1.82-3.07) | 2.04 (1.56-2.68) |
| 10-20 | 940 (2.4) | 23 (7.3) | 4.68 (3.00-7.32) | 3.83 (2.43-6.03) | 2.98 (1.86-4.78) |
| 20+ | 623 (1.6) | 14 (4.4) | 4.31 (2.48-7.49) | 3.39 (1.95-5.91) | 2.46 (1.39-4.37) |
| **Number of actinic lesions**<br>**burnt/frozen off** |  |  |  |  |  |
| None | 17,309 (44.9) | 70 (22.2) | Reference | Reference | Reference |
| 1-5 | 10,428 (27.1) | 81 (25.6) | 1.94 (1.41-2.68) | 1.84 (1.33-2.53) | 1.73 (1.25-2.39) |
| 6-10 | 3669 (9.5) | 40 (12.7) | 2.72 (1.84-4.01) | 2.43 (1.64-3.61) | 2.11 (1.42-3.14) |
| 11-20 | 2961 (7.7) | 41 (13.0) | 3.45 (2.34-5.07) | 2.98 (2.02-4.41) | 2.50 (1.68-3.70) |
| 21-50 | 2261 (5.9) | 43 (13.6) | 4.70 (3.21-6.88) | 3.99 (2.71-5.86) | 3.15 (2.12-4.67) |
| 50+ | 1888 (4.9) | 41 (13.0) | 5.37 (3.65-7.90) | 4.37 (2.95-6.48) | 3.16 (2.07-4.82) |

<sup>1</sup>Model 1- additionally adjusted for hair colour, tanning ability

### Sensitivity analyses

Censoring *in situ* melanomas occurring during follow-up made no material difference to the estimates (Table S1), nor did treating *in situ* melanoma occurring during follow-up a time-varying covariate (Table S2).

### Gene/environment interaction

The characteristics of the sub-group with a PRS were similar to the full cohort in terms of age, sex, country of birth, phenotype and history of sunburns, but participants with GWAS data were more likely to be university educated and have private health insurance, to have a history of skin cancers/AKs and to have reported undergoing physician skin checks (Table S3). Participants at highest genetic risk for melanoma (PRS tertile 3) had three-fold higher risk of melanoma than those in tertile 1 [HR T3 vs T1 3.02 (95% CI 2.01-4.53); T2 vs T1 1.50 (95% CI 0.96-2.37); Table 5]. We found a significant interaction between genetic risk and country of birth; PRS was significantly associated with invasive melanoma among people with a high (T3) but not low (T1-2) PRS (P_int_ 0.03). In contrast, we found that past history of actinic lesions was more strongly associated with invasive melanoma among people at lowest genetic risk (T1 vs T2/3 P_int_ 0.03). There was no significant difference in the association between sunburns as a youth, sunbed use or past history of skin cancers across PRS groups.

**Table 5.** Measures of UV exposure and risk of first invasive melanoma according to polygenic risk score (n=15,373).

| Variables | T1<sup>1</sup> |  | T2<sup>1</sup> |  | T3<sup>1</sup> |  | P<sup>2</sup> |
| --- | --- | --- | --- | --- | --- | --- | --- |
|  | Crude | Age & sex adj | Crude | Age & sex adj | Crude | Age & sex adj |  |
|  | HR (95%CI) | HR (95%CI) | HR (95%CI) | HR (95%CI) | HR (95%CI) | HR (95%CI) |  |
| **Born in Australia** |  |  |  |  |  |  |  |
| No | Reference | Reference | Reference | Reference | Reference | Reference |  |
| Yes | 0.76 (0.33-1.77) | 0.88 (0.38-2.04) | 0.87 (0.42-1.80) | 0.86 (0.41-1.78) | 3.01 (1.32-6.86) | 3.16 (1.39-7.22) | 0.03 |
| **Sunburns as a youth (10-20 years old)** |  |  |  |  |  |  |  |
| Never | Reference | Reference | Reference | Reference | Reference | Reference |  |
| 1-5 times | 1.27 (0.17-9.55) | 1.54 (0.20-11.73) | 2.35 (0.32-17.40) | 2.40 (0.33-17.40) | 1.10 (0.26-4.62) | 1.16 (0.28-4.85) |  |
| 6+ times | 1.37 (0.19-10.12) | 1.83 (0.25-13.49) | 1.53 (0.20-11.43) | 1.51 (0.20-11.16) | 1.49 (0.36-6.13) | 1.66 (0.41-6.73) | 0.37 |
| **Sunbed use** |  |  |  |  |  |  |  |
| Never | Reference | Reference | Reference | Reference | Reference | Reference |  |
| Ever | 1.74 (0.67-4.53) | 2.32 (0.92-5.82) | 1.30 (0.55-3.07) | 1.63 (0.69-3.86) | 0.65 (0.28-1.48) | 0.77 (0.34-1.74) | 0.29 |
| **Number of skin cancers excised** |  |  |  |  |  |  |  |
| None | Reference | Reference | Reference | Reference | Reference | Reference |  |
| 1 | 1.87 (0.59-5.97) | 1.68 (0.52-5.42) | 2.83 (1.27-6.30) | 2.79 (1.26-6.16) | 1.24 (0.65-2.38) | 1.16 (0.60-2.22) |  |
| 2+ | 4.40 (2.01-9.60) | 3.72 (1.66-8.33) | 3.06 (1.59-5.90) | 2.79 (1.47-5.29) | 2.05 (1.33-3.15) | 1.85 (1.19-2.88) | 0.28 |
| **Number of actinic lesions burnt/frozen off** |  |  |  |  |  |  |  |
| None | Reference | Reference | Reference | Reference | Reference | Reference |  |
| 1-5 | 1.30 (0.35-4.84) | 1.26 (0.33-4.81) | 1.27 (0.59-2.75) | 1.24 (0.59-2.62) | 1.74 (0.96-3.12) | 1.61 (0.89-2.90) |  |
| 6+ | 7.47 (2.83-19.72) | 6.74 (2.41-18.81) | 1.75 (0.89-3.45) | 1.55 (0.79-3.04) | 2.28 (1.34-3.87) | 1.97 (1.14-3.40) | 0.03 |

<sup>1</sup>T – tertile; <sup>2</sup>P – P-value for interaction

### Body Site

We found no evidence that risks of melanoma associated with cumulative sun exposure, average number of hours in the sun on weekdays or weekend days, or sunbed use varied across different body sites (Table S4). Being native born was significantly associated with risk of invasive melanoma of the lower limbs, but not of other body sites. The inverse association with the age at which participants moved to Australia (for those born overseas) was strongest for melanoma of the lower limbs. For sunburns, we found a high number of childhood burns was significantly associated with melanoma of the upper limbs only, while sunburns in youth and adulthood were both associated with melanoma of the trunk and upper limbs, but not other sites. A past history of keratinocyte cancer and of non-surgical treatments for other actinic lesions was associated with melanoma of all sites; the relationship was strongest for melanoma of the upper limbs.

### Sex

The associations between sun exposure variables and invasive melanoma did not differ significantly by sex (Table S5). Sunbed use was associated with melanoma in women but not men, but the difference was not statistically significant.

### All melanoma (invasive + *in situ*)

The magnitude of associations for ‘all melanoma’ (i.e. invasive + *in situ*) was attenuated for most sun exposure variables when compared with ‘invasive only’ melanomas, except for numbers of actinic lesions, which was similar (Table S6).

## DISCUSSION

We have reported risk estimates for melanoma associated with measures of lifetime ambient and personal UV exposure, and markers of cumulative UV damage in a large population-based prospective study of men and women living in a region with high ambient exposure. People who were born in Australia or migrated to Australia at young age, and those who had many sunburns all had significantly increased risks of melanoma. Past history of keratinocyte cancer and other actinic lesions were also both strongly associated with melanoma. Other measures of continuous and intermittent patterns of sun exposure were not significantly associated with melanoma in this study. We examined differences according to sex, provided estimates by body site of melanoma, and we considered both *in situ* and invasive melanoma. We also found that some associations with measures of sun exposure varied according to polygenic risk score.

The findings are important given that most of the literature on the association between sun exposure and melanoma derives from case-control studies. These studies are particularly prone to recall bias whereby cases systematically report their past exposure differently from controls on account of awareness of their diagnosis. Such bias is impossible to eradicate through statistical analysis and can only be avoided by using prospective designs. In the current study, all measures of sun exposure and other comprehensive risk factor information were collected at baseline.

Migration studies generally indicate a higher melanoma risk in individuals who spent their childhood in regions with high ambient UV radiation, and decreasing risk with older age at arrival in such regions (Holman and Armstrong, 1984; English *et al.*, 1997; Whiteman *et al.*, 2001). Our findings are consistent with those earlier reports, supporting the notion that sun exposure during the ‘critical period’ of early life is important for future melanoma development. Sunburns in childhood are often reported as posing the greatest risk for melanoma (Whiteman and Green, 1994; Gandini *et al.*, 2005). Our data suggest that a high number of sunburns increases the risk of melanoma regardless of when they are received. A proportion of the effect of sunburns in youth was mediated via nevus density, consistent with a causal association between sun exposure in early life and the development of nevi (Whiteman *et al.*, 2005).

We found no association between self-reported cumulative sun exposure and melanoma. The only other prospective study to measure cumulative/continuous sun exposure (other than “sunny vacations” or “wearing a bathing suit” (Han *et al.*, 2006; Nielsen *et al.*, 2012; Wu *et al.*, 2016)) is the E3N study (Savoye *et al.*, 2018), which also reported no association. This lack of effect with self-reported measures of continuous sun exposure is notable, especially given the strong and significant associations with objective markers of cumulative actinic damage such as numbers of excisions or treatments for skin lesions. Non-differential exposure misclassification is arguably the most likely explanation, given the modest repeatability of self-reported measures of personal sun exposure (Morze *et al.*, 2012), although other explanations (narrow range of exposure in Queensland; poor proxy for actual UV dose at target cells) are also likely.

The divergent pathway hypothesis for melanoma posits a model whereby individuals with high genetic propensity only require sun exposure to initiate melanomagenesis, whereas those with low genetic propensity require continued high levels of sun exposure to drive tumor development (Whiteman *et al.*, 1998; Whiteman *et al.*, 2003). We found some evidence for gene-environment interactions consistent with the divergent pathway hypothesis. In particular, the observation that a history of treatment for actinic lesions (a proxy for high cumulative sun exposure) was more strongly associated with melanoma among people with low than high polygenic risk accords with this hypothesis. Replicating these analyses in prospective datasets from other settings would provide a stronger test for this hypothesis.

Apart from the prospective design and large sample size, strengths of our study include the population-based sampling frame, and complete ascertainment of melanoma events during follow-up. Our analyses stratified by PRS is also novel for a prospective study of melanoma. A weakness was the relatively small number of cases, which resulted in limited power to examine differences in exposure effects on melanoma of different body sites. While our measures of past sun exposure were self-reported, most showed moderate-to-good agreement (weighted kappa 0.4-0.6) while sunbed use, history of skin cancer excisions and non-surgical treatments for actinic lesions were highly reproducible (weighted kappas>0.8) (Morze *et al.*, 2012). We did not confirm the histology of self-reported skin cancers that had been excised (except for melanomas, which were excluded). Histologically confirmed incident KCs are an endpoint of the QSkin study, however, and greater duration of follow-up will enable examination of these relationships according to KC type. Lastly, we are aware of some selection forces that may have contributed to the findings in unpredictable ways. Study participants, and those with genotypic data, were more highly educated and were more likely to have had a history of skin cancer than the general population, which may also limit generalizability.

In summary, we have reported estimates of the risk of melanoma associated with measures of sun exposure from a large prospective study conducted in a setting of high ambient insolation. We found some evidence of gene-environment interactions that are consistent with divergent pathways to melanoma development. In clinical practice, the advent of genomic medicine will likely have implications for patient care, and clinicians may need to consider genetic risk and its interaction with environmental exposures. In this context, our findings may help to identify and inform persons at high risk of melanoma (over and above the role of phenotype) who stand to benefit most from adopting sun protective behaviors.

## METHODS

### Study population

The QSkin Sun and Health Study is a prospective cohort study of men and women aged 40-69 years, randomly sampled from the Queensland population (n = 43,794) in 2011. The study design and characteristics of the cohort have been published (Olsen *et al.*, 2012), and the baseline survey is available online: https://qskin.qimrberghofer.edu.au/page/About/Baseline_survey. The survey included questions about sun exposure and sun protection, demographic items, pigmentary and phenotypic characteristics, family history of melanoma, past history of skin cancer and general medical history. The repeatability and validity of these items has been reported previously (Morze *et al.*, 2012).

We restricted our analyses to participants who reported white European ancestry and excluded those with a prior history of invasive or *in situ* melanoma (n=1,831); the final cohort eligible for these analyses included 38,833 participants.

A polygenic risk score (PRS) for melanoma was calculated for a sub-set of this cohort (n=15,373; 39.6%) who provided a DNA sample (see methods below).

The Human Research Ethics Committee at the QIMR Berghofer Medical Research Institute approved the study, and all participants gave written informed consent to take part.

### Outcomes

We examined two separate outcomes: 1) invasive; and 2) invasive + *in situ* melanomas. Notifications for melanoma are mandatory by law in Queensland, and data on all melanoma diagnoses from baseline up to 31 December 2015 were obtained from the Queensland Cancer Registry, supplemented by pathology reports from major pathology companies servicing Queensland.

### Exposure assessment

We considered three groups of UV radiation exposure variables: (1) ambient sun exposure (country of birth: Australia or elsewhere; latitude of birth; region of residence; number of years lived outside Australia); (2) personal exposure to UV radiation (number of sunburns in childhood, adolescence and adulthood; cumulative sun exposure; average number of hours in the sun on weekdays and weekend days; sunbed use [indoor tanning]); and (3) proxies of high cumulative UV radiation exposure [self-reported history of skin cancers (not melanomas) excised surgically; actinic skin lesions treated destructively].

Participants were asked to report their country of birth, the age they moved permanently to Australia, years of life lived in three regions of Australia (as depicted by a map and labelled ‘Northern’, ‘Central’ and ‘Southern’), and the region in which they lived the longest as a child/youth (up to age 20 years). The survey asked about the number of times the participant had been sunburned ‘so badly that you were sore for at least 2 days, or your skin peeled’ as a child, as a teenager/youth, and as an adult (possible responses were ‘never’, ‘1-5 times’, ‘6-10 times’, ‘11-20 times’, ‘21-50 times’ and ‘50+ times’) and about the number of times they had used sunbeds. Participants were also asked to report the number of hours ‘typically spent outdoors and in the sun each day’ at ages 10-19, 20-29, 30-39 years and in the past year, separately for week days and weekend days. We calculated a measure of cumulative exposure, and an average of hours spent outdoors on weekdays and weekend days, all expressed in quartiles. The survey asked about sunscreen and hat use ‘when outside in the sun during the past year’. Sunbed use, self-reported history of skin cancer excisions and non-surgical treatments for actinic lesions were highly reproducible (weighted kappas>0.8). Other measures of sun exposure showed moderate-to-good agreement (Morze *et al.*, 2012).

### Genotyping, imputation and data quality control

A total of 17,965 QSkin participants were genotyped using the Illumina Global Screening array (San Diego, CA, USA). Genotype data was cleaned using Illumina GenomeStudio/BeadStudio (San Diego, CA, USA) and PLINK (v1.9) (Chang *et al.*, 2015). We excluded participants with > 5% genotype missingness (n=322), those who were related to another sample at identity by descent 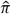 score > 0.1875 [i.e. closer than a 2^nd^ degree cousin (n=400)], or who were outliers from European reference populations (> 6 SD on PC1 and PC2, n=378) (final n=16,687 as a related pair were also population outliers). After removing 198,387 SNPs with GenTrain score <0.6, Hardy-Weinberg P-value <1 × 10^-6^, or a minor allele frequency (MAF) <1%, the remaining 496,695 SNPs were imputed to the Haplotype Reference Consortium v1.1 panel (McCarthy *et al.*, 2016) using the University of Michigan Imputation Server. We performed genotype phasing with Eagle 2 (Loh *et al.*, 2016) and genotype imputation by minimac version 3 (Das *et al.*, 2016). The resulting imputed GWAS data was analysed as dosage data filtered to imputation quality score 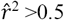 and MAF>0.001.

### Polygenic risk score

We calculated a PRS for 15,373 of the 38,832 participants who were eligible for these analyses (39.6%) using summary statistics from a melanoma GWAS meta-analysis for 12,874 cases and 23,203 controls (Law *et al.*, 2015); full details are provided in Supplementary Methods. The PRS was categorised into tertiles based on the distribution of the analysis sample.

### Imputation of missing data

Missing values for sun exposure variables ranged from <1% to 10%; the sunburn variables had the highest amount of missing data (9.7%, 3.4% and 4.7% for sunburns in childhood, adolescence and adulthood, respectively). To avoid losing observations due to missing covariate data during model development, we imputed missing values using PROC MI in SAS v9.4 (SAS Institute, Cary, NC), assuming that data were missing at random. We included all sun exposure variables and the outcome variable (Moons *et al.*, 2006) in the imputation step; imputation was run over 50 cycles to generate 50 data sets.

### Statistical analysis

We used Cox proportional hazards models to estimate the total effect of each measure of sun exposure on the risk of first incident melanoma while taking account of the sociodemographic and phenotypic factors as well as sun protection practices. Choice of covariates was guided by direct acyclic graphs and *a priori* knowledge. We first examined each factor unadjusted, and then adjusted for age and sex. We then sequentially considered covariates from three groups: (1) those related to pigmentation; (2) family history of melanoma; and (3) sun protection behaviors. For sun exposure variables related to ambient UV levels we considered sun protection behaviors only. For proxies of high cumulative UV exposure we considered pigmentary factors only. Inclusion of covariates in groups 2 and 3 did not result in material change to the estimates of effect and thus models with these factors are not presented.

For analyses of the primary outcome (invasive melanoma) we ignored all *in situ* melanomas diagnosed during follow-up. We conducted sensitivity analyses to examine the influence of this approach firstly by censoring the *in situ* melanoma cases, and secondly by treating *in situ* cases occurring during follow-up as a time-varying covariate in the model. For our primary outcome we examined the associations for melanoma of different body sites (trunk, head and neck, upper limbs, lower limbs) and conducted analyses stratified by sex and PRS tertiles; interaction was assessed using cross-product terms.

Finally, sunburn events in early life are associated with nevus density (Whiteman *et al.*, 2005), which influences melanoma risk (Olsen *et al.*, 2010), so we examined the degree to which nevus density accounted for the influence of sunburns in childhood and youth on melanoma risk by conducting mediation analyses using the method of Valeri and Vanderweele (Valeri and Vanderweele, 2013).

All models were adjusted for death as a competing risk (Fine and Gray, 1999), and statistical significance was inferred at *P* < .05. All analyses were conducted using SAS 9.4 software (SAS Institute, Cary, NC).

## ABBREVIATIONS

UV: ultraviolet
BCC: basal cell carcinoma
SCC: squamous cell carcinoma
KC: keratinocyte cancer
PRS: polygenic risk score
GWAS: genome-wide association study
MAF: minor allele frequency
HR: hazard ratio
SD: standard deviation

## ACKNOWLEDGEMENTS

This study was supported in part by the National Health and Medical Research Council (NHMRC) of Australia (grant number 552429). DCW, REN and SM are supported by fellowships from the NHMRC. This work was conducted using the UK Biobank Resource (application number 25331).

