## Supplementary methods for "Ultraviolet radiation exposure and melanoma: evidence for gene-environment interaction in a large prospective cohort"

### Calculation of polygenic risk score

We calculated a polygenic risk score (PRS) for 15,373 participants who were eligible for these analyses (39.6%) and had GWAS data available. The PRS was calculated using summary statistics from a melanoma GWAS meta-analysis for 12,874 cases and 23,203 controls.[1] Consistent with the meta-analysis,[1] the PRS included only SNPs with an imputation quality score (RSQ or INFO) 0.8, MAF > 0.001 and present in 2 or more studies; the QSkin cohort was not included in the meta-analysis.

Linkage Disequilibrium (LD) between the meta-analysis SNPs was determined using a reference panel generated from 5,000 White British individuals from the UK Biobank (Application ID 25331). UK Biobank samples' HRC v1 imputed data was filtered to those with an imputation quality RSQ > 0.3, a Hardy-Weinberg P-value > $1 \times 10^{-6}$ and missingness < 0.03, and then converted to best guess genotypes.

Using the intersection of the available, cleaned SNPs for the QSkin GWAS data, the melanoma meta-analysis, and the UK Biobank LD panel, this overlapping set of SNPs was pruned (LD $r^2$ threshold < 0.025 over a 10 megabase window) to 123,038 independent SNPs unselected for melanoma P-value using PLINK v1.90b6.6.[2] Due to the strong cutaneous melanoma association at the *MC1R* on chromosome 16, even at the very strict LD $r^2$ threshold of 0.025 multiple additional SNPs were present post pruning near *MC1R*, each with LD $r^2$ 0.01-0.02 with *MC1R* red hair alleles. To address this, the terminal 5 megabases of chromosome 16 was removed from the pruned set of SNPs, and instead the 7 *MC1R* missense alleles passing QC across all sets (those strongly associated with melanoma, red hair and pigmentation[3] termed 'R' alleles rs1805006 ASP84GLU, rs1805007 ARG151CYS, rs1110400 I155T, rs1805008 ARG160TRP, and less strongly associated 'r' allele rs1805005 VAL60LEU, rs2228479 VAL92MET, rs885479 R163Q were retained for this region.

PRS scores were generated by the --score function of PLINK v1.9, which sums the log(OR) of the fixed effects meta-analysis for each copy of the effect allele, with the non-effect allele scored 0. To determine the most powerful set of SNPs 10 scores were constructed using different fixed effects meta-analysis p-value bins (S1 0–5 × $10^{-8}$; S2 0–5 × $10^{-7}$; S3 0–5 × $10^{-6}$; S4 0–5 × $10^{-5}$; S5 0 5 × $10^{-4}$; S6 0–5 × $10^{-3}$; S7 0–0.01; S8 0–0.05; S9 0–0.10; S10 0–1). PRS P-value bins were assessed by their goodness of fit using nagelkerke psuedo-R2 when regressed onto QSkin melanoma status using logistic regression together with the first 10 principal components from the PCA analysis used to exclude population outliers in R v3.5.1.[4] Bin 2 ($0 < P < 5 \times 10^{-7}$) achieved the highest goodness of fit, and included 28 independent SNPs (Appendix 1).

**Appendix 1**. SNPs used in calculation of polygenic risk for melanoma.

| CHR | BP | SNP | EA | NEA | BETA | P | AF_HRC |
|---|---|---|---|---|---|---|---|
| 1 | 150856153 | rs12410869 | T | G | -0.13045 | 5.21E-013 | 0.380259 |
| 1 | 226608104 | rs1858550 | A | C | -0.142601 | 1.68E-013 | 0.370034 |
| 2 | 38276549 | rs6750047 | G | A | -0.0883942 | 2.92E-007 | 0.541099 |
| 2 | 202176294 | rs7582362 | G | A | -0.113169 | 8.88E-009 | 0.719002 |
| 5 | 1320247 | rs380286 | A | G | 0.151519 | 1.66E-017 | 0.446628 |
| 5 | 33952378 | rs250417 | C | G | 0.890645 | 2.30E-012 | 0.926748 |
| 6 | 21163919 | rs6914598 | C | T | 0.108226 | 2.56E-008 | 0.317324 |
| 7 | 16984280 | rs1636744 | T | C | 0.105441 | 1.84E-009 | 0.382676 |
| 7 | 130754812 | rs4731742 | C | G | 0.11805 | 4.29E-007 | 0.752587 |
| 8 | 37482669 | rs56122426 | A | G | -0.156186 | 2.04E-007 | 0.160132 |
| 9 | 21803183 | rs935055 | C | G | -0.213812 | 2.65E-032 | 0.523576 |
| 9 | 22060833 | rs77283072 | A | G | 0.136714 | 3.75E-009 | 0.171928 |
| 9 | 109060830 | rs10739221 | C | T | -0.119798 | 9.58E-009 | 0.740838 |
| 9 | 110698716 | rs7041168 | T | G | -0.0957403 | 1.31E-007 | 0.512766 |
| 11 | 68919649 | rs2290419 | G | A | -0.201382 | 4.20E-007 | 0.0576378 |
| 11 | 69367118 | rs498136 | C | A | -0.115748 | 1.01E-010 | 0.640407 |
| 11 | 89028043 | rs10830253 | G | T | 0.194003 | 1.01E-026 | 0.295981 |
| 11 | 108187689 | rs73008229 | A | G | -0.188259 | 1.38E-012 | 0.144472 |
| 12 | 13075202 | rs2111398 | G | A | 0.097036 | 3.68E-007 | 0.437512 |
| 14 | 91185865 | rs184628474 | A | G | -1.03282 | 4.63E-014 | 0.03919 |
| 15 | 28335820 | rs4778138 | G | A | -0.177692 | 3.11E-009 | 0.180582 |
| 16 | 54115829 | rs12596638 | A | G | 0.142887 | 1.81E-009 | 0.178596 |
| 16 | 68796746 | rs35158985 | G | A | 0.101383 | 1.31E-007 | 0.327903 |
| 16 | 89986117 | rs1805007 | T | C | 0.615186 | 4.68E-083 | 0.0634586 |
| 16 | 89986144 | rs1805008 | T | C | 0.316707 | 6.26E-025 | 0.0663536 |
| 20 | 32538391 | rs62211989 | C | G | 0.350446 | 1.63E-030 | 0.0576994 |
| 21 | 42743496 | rs408825 | T | C | 0.140718 | 3.21E-015 | 0.586834 |
| 22 | 38545942 | rs132941 | C | T | -0.123072 | 1.61E-012 | 0.451078 |

Melanoma risk effect size from a melanoma GWAS meta-analysis,[1] and are the log of the odds ratio (BETA) for the effect allele (EA). For reference we also report the hg19 chromosome (CHR) and base pair (BP) position, the non-effect allele (NEA), the fixed effects meta-analysis p-value (P), and the effect allele frequency from the Haplotype Reference Consortium.[5]

**Table S1.** Measures of **s**un exposure and risk of invasive melanoma (*in situ* melanomas occurring during follow-up are censored).

| | Invasive melanoma | | |
|---|---|---|---|
| | **No** (n=38,529) | **Yes** (n=303) | |
| **Variables** | **n (%)** | **n (%)** | **HR (95%CI)**[1] |
| **Born in Australia** | | | |
| No | 6783 (17.6) | 40 (13.2) | Reference |
| Yes | 31,746 (82.4) | 263 (86.8) | 1.50 (1.08-2.10) |
| **Latitude of birth** | | | |
| ≥45º S/N | 4504 (11.7) | 25 (8.3) | Reference |
| <45º S/N | 34,025 (88.3) | 278 (91.8) | 1.62 (1.07-2.43) |
| **Region of Australia lived longest up to age 20 years** | | | |
| Southern region | 11,651 (33.7) | 96 (33.9) | Reference |
| Central region | 17,039 (49.4) | 153 (54.1) | 1.15 (0.89-1.48) |
| Northern region | 5816 (16.8) | 34 (12.0) | 0.76 (0.51-1.12) |
| **Region of Australia lived longest over lifetime** | | | |
| Southern region | 8779 (22.8) | 71 (23.4) | Reference |
| Central region | 22,907 (55.5) | 194 (64.0) | 1.06 (0.81-1.39) |
| Northern region | 6843 (17.8) | 38 (12.5) | 0.70 (0.47-1.03) |
| **Age moved to Australia** | | | |
| Born in Australia | 31,786 (82.5) | 263 (86.8) | Reference |
| 1-10 years | 1875 (4.9) | 11 (3.6) | 0.70 (0.38-1.27) |
| 11-20 years | 1092 (2.8) | 12 (4.0) | 1.24 (0.69-2.21) |
| 20+ years | 3776 (9.8) | 17 (5.6) | 0.50 (0.30-0.81) |
| **Sunburns as a child (less than 10 years)** | | | |
| Never | 8082 (21.0) | 57 (18.8) | Reference |
| 1-5 times | 17,588 (45.6) | 143 (47.2) | 1.02 (0.74-1.40) |
| 6-10 times | 6733 (17.5) | 53 (17.5) | 0.92 (0.62-1.38) |
| 11-20 times | 3643 (9.5) | 22 (7.3) | 0.79 (0.47-1.31) |
| 21-50 times | 1748 (4.5) | 15 (5.0) | 1.01 (0.57-1.81) |
| >50 times | 735 (1.9) | 13 (4.3) | 1.83 (0.96-3.48) |
| **Sunburns as a youth (10-20 years old)** | | | |
| Never | 1788 (4.6) | 9 (3.0) | Reference |
| 1-5 times | 16,799 (43.6) | 127 (41.9) | 1.55 (0.77-3.14) |
| 6-10 times | 9582 (24.9) | 80 (26.4) | 1.73 (0.85-3.55) |
| 11-20 times | 6420 (16.7) | 49 (16.2) | 1.55 (0.74-3.23) |
| 21-50 times | 2887 (7.5) | 20 (6.6) | 1.42 (0.63-3.21) |
| >50 times | 1053 (2.7) | 18 (5.9) | 3.13 (1.35-7.25) |
| **Sunburns as an adult (more than 20 years old)** | | | |
| Never | 6231 (16.2) | 40 (13.2) | Reference |
| 1-5 times | 20,465 (53.1) | 160 (52.8) | 1.39 (0.97-1.99) |
| 6-10 times | 6398 (16.6) | 55 (18.2) | 1.50 (0.98-2.28) |
| 11-20 times | 3415 (8.9) | 20 (6.6) | 1.03 (0.59-1.77) |
| 21-50 times | 1462 (3.8) | 22 (7.3) | 2.45 (1.44-4.17) |

| | Invasive melanoma | | |
|---|---|---|---|
| | **No** (n=38,529) | **Yes** (n=303) | |
| **Variables** | **n (%)** | **n (%)** | **HR (95%CI)**[1] |
| >50 times | 558 (1.5) | 6 (2.0) | 1.70 (0.71-4.06) |
| **Cumulative sun exposure**[2] | | | |
| Q1 | 9344 (24.3) | 69 (22.8) | Reference |
| Q2 | 9676 (25.1) | 66 (21.8) | 0.87 (0.62-1.23) |
| Q3 | 9729 (25.3) | 84 (27.7) | 0.97 (0.69-1.35) |
| Q4 | 9781 (25.4) | 84 (27.7) | 0.83 (0.59-1.18) |
| **Average number of hours in the sun on weekdays**[2] | | | |
| Q1 | 9033 (23.4) | 71 (23.4) | Reference |
| Q2 | 9446 (24.5) | 73 (24.1) | 1.00 (0.71-1.40) |
| Q3 | 10,156 (26.4) | 80 (26.4) | 0.93 (0.67-1.29) |
| Q4 | 9894 (25.7) | 79 (26.1) | 0.86 (0.61-1.20) |
| **Average number of hours in the sun on weekend days**[2] | | | |
| Q1 | 9695 (25.2) | 76 (25.1) | Reference |
| Q2 | 9860 (25.6) | 81 (26.7) | 1.04 (0.74-1.45) |
| Q3 | 9780 (25.4) | 69 (22.8) | 0.91 (0.65-1.27) |
| Q4 | 9194 (23.9) | 77 (25.4) | 1.02 (0.72-1.44) |
| **Sunbed use** | | | |
| Never | 34,413 (89.3) | 273 (90.1) | Reference |
| Ever | 4116 (10.7) | 30 (9.9) | 1.31 (0.89-1.92) |
| **Number of skin cancers excised** | | | |
| None | 23,540 (61.1) | 118 (38.9) | Reference |
| 1 | 5368 (13.9) | 45 (14.9) | 1.48 (1.04-2.10) |
| 2-10 | 8055 (20.9) | 106 (25.0) | 2.03 (1.54-2.68) |
| 10-20 | 942 (2.4) | 21 (6.9) | 2.86 (1.75-4.67) |
| 20+ | 624 (1.6) | 13 (4.3) | 2.40 (1.33-4.34) |
| **Number of actinic lesions burnt/frozen off** | | | |
| None | 17,311 (44.9) | 68 (22.4) | Reference |
| 1-5 | 10,429 (27.1) | 80 (26.4) | 1.76 (1.27-2.43) |
| 6-10 | 3672 (9.5) | 37 (12.2) | 2.02 (1.34-3.05) |
| 11-20 | 2963 (7.7) | 39 (12.9) | 2.46 (1.64-3.68) |
| 21-50 | 2263 (5.9) | 41 (13.5) | 3.10 (2.08-4.64) |
| 50+ | 1891 (4.9) | 38 (12.5) | 3.05 (1.97-4.71) |

[1]Born in Australia, latitude of birth, region of Australia lived longest (as a child; lifetime) adjusted for age and sex.
All other variables adjusted for age, sex, hair colour and tanning ability.
[2]Q - Quartile

**Table S2.** Measures of **s**un exposure and risk of invasive melanoma (*in situ* melanomas treated as a time-varying covariate).

| | Invasive melanoma | | |
|---|---|---|---|
| | **No** (n=39,816) | **Yes** (n=346) | |
| **Variables** | **n (%)** | **n (%)** | **HR (95%CI)**[1] |
| **Born in Australia** | | | |
| No | 6942 (17.5) | 43 (12.4) | Reference |
| Yes | 32,836 (82.6) | 303 (87.6) | 1.59 (1.16-2.20) |
| **Latitude of birth** | | | |
| ≥45º S/N | 4591 (11.5) | 27 (7.8) | Reference |
| <45º S/N | 35,174 (88.5) | 319 (92.2) | 1.69 (1.14-2.51) |
| **Region of Australia lived longest up to age 20 years** | | | |
| Southern region | 11,773 (34.2) | 105 (33.1) | Reference |
| Central region | 16,944 (49.1) | 173 (54.6) | 1.19 (0.93-1.51) |
| Northern region | 5757 (16.7) | 39 (12.3) | 0.80 (0.56-1.16) |
| **Region of Australia lived longest over lifetime** | | | |
| Southern region | 8602 (22.2) | 74 (21.9) | Reference |
| Central region | 23,206 (60.0) | 218 (64.5) | 1.12 (0.86-1.45) |
| Northern region | 6905 (17.8) | 46 (13.6) | 0.79 (0.54-1.14) |
| **Age moved to Australia** | | | |
| Born in Australia | 32,836 (82.8) | 303 (87.6) | Reference |
| 1-10 years | 1877 (4.7) | 13 (3.8) | 0.74 (0.42-1.28) |
| 11-20 years | 1111 (2.8) | 12 (3.5) | 1.10 (0.62-1.96) |
| 20+ years | 3834 (9.7) | 18 (5.2) | 0.47 (0.29-0.75) |
| **Sunburns as a child (less than 10 years)** | | | |
| Never | 7417 (20.6) | 57 (18.2) | Reference |
| 1-5 times | 16,401 (45.5) | 147 (46.8) | 1.10 (0.82-1.51) |
| 6-10 times | 6445 (17.9) | 52 (16.6) | 1.00 (0.68-1.47) |
| 11-20 times | 3551 (9.9) | 30 (9.6) | 1.02 (0.65-1.61) |
| 21-50 times | 1594 (4.4) | 17 (5.4) | 1.23 (0.71-2.15) |
| >50 times | 1058 (2.8) | 11 (3.5) | 1.67 (0.84-3.32) |
| **Sunburns as a youth (10-20 years old)** | | | |
| Never | 1679 (4.4) | 8 (3.4) | Reference |
| 1-5 times | 16,677 (43.3) | 144 (42.9) | 1.80 (0.88-3.67) |
| 6-10 times | 9671 (25.1) | 83 (24.7) | 1.85 (0.89-3.84) |
| 11-20 times | 6513 (16.9) | 56 (16.7) | 1.86 (0.88-3.91) |
| 21-50 times | 2895 (7.5) | 24 (7.1) | 1.73 (0.77-3.90) |
| >50 times | 1054 (2.7) | 21 (6.3) | 4.19 (1.84-9.55) |
| **Sunburns as an adult (more than 20 years old)**[2] | | | |
| Never | 6086 (16.0) | 44 (13.3) | Reference |
| 1-5 times | 20,106 (52.9) | 166 (50.0) | 1.28 (0.91-1.79) |
| 6-10 times | 6415 (16.9) | 61 (18.4) | 1.55 (1.05-2.29) |
| 11-20 times | 3446 (9.1) | 30 (9.0) | 1.41 (0.89-2.25) |
| 21-50 times | 1433 (3.8) | 24 (7.2) | 2.65 (1.61-4.38) |

| | Invasive melanoma | | |
|---|---|---|---|
| | **No** (n=39,816) | **Yes** (n=346) | |
| **Variables** | **n (%)** | **n (%)** | **HR (95%CI)**[1] |
| >50 times | 537 (1.4) | 7 (2.1) | 2.07 (0.93-4.62) |
| **Cumulative sun exposure** | | | |
| Q1 | 8495 (24.7) | 68 (23.6) | Reference |
| Q2 | 8641 (25.1) | 60 (20.8) | 0.88 (0.62-1.24) |
| Q3 | 8582 (25.0) | 72 (25.0) | 0.99 (0.70-1.39) |
| Q4 | 8659 (25.2) | 88 (30.6) | 1.16 (0.84-1.61) |
| **Average number of hours in the sun on weekdays**[2] | | | |
| Q1 | 8500 (23.8) | 68 (23.1) | Reference |
| Q2 | 8775 (24.6) | 74 (25.1) | 1.09 (0.78-1.52) |
| Q3 | 9389 (26.3) | 72 (24.4) | 0.99 (0.71-1.38) |
| Q4 | 9055 (25.4) | 81 (27.5) | 1.19 (0.86-1.65) |
| **Average number of hours in the sun on weekend days**[2] | | | |
| Q1 | 8853 (24.7) | 74 (24.3) | Reference |
| Q2 | 9092 (25.4) | 73 (23.9) | 1.07 (0.77-1.48) |
| Q3 | 9223 (25.7) | 81 (26.6) | 1.23 (0.90-1.69) |
| Q4 | 8699 (24.3) | 77 (25.3) | 1.32 (0.95-1.82) |
| **Sunbed use** | | | |
| Never | 35,360 (89.4) | 312 (90.7) | Reference |
| Ever | 4200 (10.6) | 32 (9.3) | 1.12 (0.78-1.62) |
| **Number of skin cancers excised** | | | |
| None | 23,637 (59.8) | 123 (35.8) | Reference |
| 1 | 5607 (14.2) | 51 (14.8) | 1.57 (1.13-2.19) |
| 2-10 | 8565 (21.7) | 127 (36.9) | 2.27 (1.75-2.95) |
| 10-20 | 1029 (2.6) | 24 (7.0) | 2.91 (1.81-4.69) |
| 20+ | 677 (1.7) | 19 (5.5) | 3.34 (2.00-5.59) |
| **Number of actinic lesions burnt/frozen off** | | | |
| None | 17,500 (44.2) | 73 (21.2) | Reference |
| 1-5 | 10,708 (27.0) | 84 (24.4) | 1.65 (1.20-2.26) |
| 6-10 | 3820 (9.6) | 44 (12.8) | 2.18 (1.48-3.20) |
| 11-20 | 3114 (7.9) | 46 (13.3) | 2.63 (1.81-3.84) |
| 21-50 | 2440 (6.2) | 48 (13.9) | 3.16 (2.16-4.64) |
| 50+ | 2032 (5.1) | 50 (14.5) | 3.64 (2.45-5.41) |

[1]Born in Australia, latitude of birth, region of Australia lived longest (as a child; lifetime) adjusted for age and sex.
All other variables adjusted for age, sex, hair colour and tanning ability.
[2]Q - Quartile

**Table S3.** Characteristics of entire cohort (n=38,832) and stratified by PRS availability.

| Characteristic | Whole cohort (n=38,832) N (%)[a] | Without PRS (n=23,459) N (%)[a] | With PRS (n=15,373) N (%)[a] | Chi-Square *P*-value |
|---|---|---|---|---|
| **Age** | | | | |
| 40-49 | 10,278 (26.5) | 6928 (29.5) | 3350 (21.8) | |
| 50-59 | 14,724 (37.9) | 8698 (37.1) | 6026 (39.2) | |
| ≥60 | 13,830 (35.6) | 7833 (33.4) | 5997 (39.0) | <0.001 |
| **Mean age (SD)** | 56.0 (8.2) | 56.9 (7.9) | 55.5 (8.3) | <0.001[1] |
| **Sex** | | | | |
| Female | 21,067 (54.3) | 12,632 (53.8) | 8435 (54.9) | |
| Male | 17,765 (45.7) | 10,827 (46.2) | 6938 (45.1) | 0.05 |
| **Education** | | | | |
| No school | 3286 (8.5) | 2313 (9.9) | 973 (6.3) | |
| School or intermediate | 6532 (16.8) | 4148 (17.7) | 2384 (15.5) | |
| High school | 7641 (19.7) | 4775 (20.4) | 2866 (18.6) | |
| Trade/apprenticeship | 4007 (10.3) | 2564 (10.9) | 1443 (9.4) | |
| Certificate/diploma | 7730 (19.9) | 4482 (19.1) | 3248 (21.1) | |
| University | 9636 (24.8) | 5177 (22.1) | 4459 (29.0) | <0.001 |
| **Private health insurance** | | | | |
| Yes | 26,176 (67.4) | 15,138 (64.5) | 11,038 (71.8) | |
| No | 12,656 (32.6) | 8321 (35.5) | 4335 (28.2) | <0.001 |
| **Native born** | | | | |
| Yes | 32,009 (82.4) | 19,302 (82.3) | 12,707 (82.7) | |
| No | 6823 (17.6) | 4157 (17.7) | 2666 (17.3) | 0.34 |
| **Skin colour** | | | | |
| Fair | 23,695 (61.0) | 13,963 (59.5) | 9732 (63.3) | |
| Medium | 12,554 (32.3) | 7777 (33.2) | 4777 (31.1) | |
| Olive/dark | 2583 (6.7) | 1719 (7.3) | 864 (5.6) | <0.001 |
| **Skin tanning ability** | | | | |
| Not tan | 2536 (6.5) | 1519 (6.5) | 1017 (6.6) | |
| Tan a little | 8187 (21.1) | 4762 (20.3) | 3425 (22.3) | |
| Tan moderately | 19,448 (50.1) | 11,752 (50.1) | 7696 (50.1) | |
| Tan deeply | 8661 (22.3) | 5426 (23.1) | 3235 (21.0) | <0.001 |
| **Hair colour** | | | | |
| Red/Auburn | 2263 (5.8) | 1361 (5.8) | 902 (5.9) | |
| Dark brown | 13,071 (33.7) | 7890 (33.6) | 5181 (33.7) | |
| Blond | 5602 (14.4) | 3456 (14.7) | 2146 (14.0) | |
| Black | 3044 (7.8) | 1854 (7.9) | 1190 (7.7) | |
| Light brown | 14,852 (38.2) | 8898 (37.9) | 5954 (38.7) | 0.22 |
| **Freckling on face at age 21** | | | | |
| None | 18,039 (46.5) | 11,122 (47.4) | 6917 (45.0) | |
| A few | 12,276 (31.6) | 7360 (31.4) | 4916 (32.0) | |
| Some | 6136 (15.8) | 3598 (15.3) | 2538 (16.5) | |
| Many | 2381 (6.1) | 1379 (5.9) | 1002 (6.5) | <0.001 |
| **Moles at age 21** | | | | |
| None | 11,100 (28.6) | 6902 (29.4) | 4198 (27.3) | |
| A few | 20,709 (53.3) | 12,338 (52.6) | 8371 (54.5) | |

| Characteristic | Whole cohort (n=38,832) N (%)[a] | Without PRS (n=23,459) N (%)[a] | With PRS (n=15,373) N (%)[a] | Chi-Square *P*-value |
|---|---|---|---|---|
| Some | 5846 (15.1) | 3546 (15.1) | 2300 (15.0) | |
| Many | 1177 (3.0) | 673 (2.9) | 504 (3.3) | <0.001 |
| **Sunburns as an adult** | | | | |
| Never | 6271 (16.1) | 3800 (16.2) | 2471 (16.1) | |
| 1-5 times | 20,625 (53.1) | 12,361 (52.7) | 8264 (53.8) | |
| 6-10 times | 6453 (16.6) | 3910 (16.7) | 2543 (16.5) | |
| 11-20 times | 3435 (8.8) | 2094 (8.9) | 1341 (8.7) | |
| 21-50 times | 1484 (3.8) | 932 (4.0) | 552 (3.6) | |
| >50 times | 564 (1.5) | 362 (1.5) | 202 (1.3) | 0.09 |
| **Number of skin cancers excised** | | | | |
| None | 23,658 (60.9) | 14,742 (62.8) | 8916 (58.0) | |
| 1 | 5413 (13.9) | 2161 (13.7) | 3252 (14.1) | |
| 2-10 | 8161 (21.0) | 4553 (19.4) | 3608 (23.5) | |
| 11-20 | 963 (2.5) | 518 (2.2) | 445 (2.9) | |
| 20+ | 637 (1.6) | 336 (1.4) | 301 (2.0) | <0.001 |
| **Number of actinic lesions burnt/frozen off** | | | | |
| None | 17,379 (44.8) | 11,252 (48.0) | 6127 (39.9) | |
| 1-5 | 10,509 (27.1) | 6270 (26.7) | 4239 (27.6) | |
| 6-10 | 3709 (9.6) | 2129 (9.1) | 1580 (10.3) | |
| 11-20 | 3002 (7.7) | 1659 (7.1) | 1343 (8.7) | |
| 21-50 | 2304 (5.9) | 1188 (5.1) | 1116 (7.3) | |
| 50+ | 1929 (5.0) | 961 (4.1) | 968 (6.3) | <0.001 |
| **Physician skin checks in past 3 years** | | | | |
| Never | 10,598 (27.3) | 6909 (29.5) | 3689 (24.0) | |
| Once | 11,798 (30.4) | 7208 (30.7) | 4590 (29.9) | |
| 2-5 times | 13,943 (35.9) | 7957 (33.9) | 5986 (38.9) | |
| More than 5 times | 2493 (6.4) | 1385 (5.9) | 1108 (7.2) | <0.001 |

[1]*P*-value for significant difference in mean values (Ryan-Einot-Gabriel-Welsch multiple range test).

**Table S4.** Measures of sun exposure and risk of first invasive melanoma, stratified by body site of melanoma[1].

| Variables | HR (95% CI)[2] | | | | P-interaction |
|---|---|---|---|---|---|
| | **Trunk** (n=110) | **Head/neck** (n=48) | **Upper limbs** (n=83) | **Lower limbs** (n=64) | |
| **Born in Australia** | | | | | |
| No | Reference | Reference | Reference | Reference | |
| Yes | 1.48 (0.86-2.54) | 1.15 (0.54-2.48) | 1.14 (0.64-2.02) | 3.27 (1.19-8.99) | 0.04 |
| **Latitude of birth** | | | | | |
| ≥45º S/N | Reference | Reference | Reference | Reference | |
| <45º S/N | 1.91 (0.93-3.92) | 1.62 (0.58-4.56) | 1.07 (0.56-2.07) | 2.04 (0.74-5.61) | 0.008 |
| **Region of Australia lived longest up to age 20 years** | | | | | |
| Southern region | Reference | Reference | Reference | Reference | |
| Central region | 0.89 (0.59-1.35) | 1.18 (0.57-2.43) | 1.45 (0.87-2.41) | 1.31 (0.77-2.26) | |
| Northern region | 0.59 (0.30-1.14) | 1.76 (0.76-4.07) | 0.96 (0.45-2.03) | 0.51 (0.19-1.36) | <0.001 |
| **Region of Australia lived longest over lifetime** | | | | | |
| Southern region | Reference | Reference | Reference | Reference | |
| Central region | 0.84 (0.55-1.30) | 1.09 (0.54-2.18) | 1.38 (0.79-2.41) | 1.30 (0.70-2.42) | |
| Northern region | 0.61 (0.32-1.15) | 0.83 (0.32-2.15) | 0.91 (0.42-1.97) | 0.69 (0.28-1.73) | <0.001 |
| **Age moved to Australia** | | | | | |
| Born in Australia | Reference | Reference | Reference | Reference | |
| 1-10 years | 0.86 (0.35-2.11) | 0.41 (0.06-3.00) | 1.45 (0.63-3.33) | - | |
| 11-20 years | 1.14 (0.42-3.11) | 1.35 (0.33-5.60) | 1.16 (0.36-3.69) | 1.40 (0.44-4.48) | |
| 20+ years | 0.47 (0.21-1.08) | 0.95 (0.37-2.46) | 0.55 (0.22-1.36) | 0.14 (0.02-0.98) | <0.001 |
| **Sunburns as a child (less than 10 years)** | | | | | |
| Never | Reference | Reference | Reference | Reference | |
| 1-5 times | 1.16 (0.66-2.03) | 1.04 (0.45-2.42) | 1.25 (0.65-2.38) | 0.89 (0.47-1.69) | |
| 6-10 times | 1.06 (0.53-2.11) | 1.18 (0.45-3.12) | 1.01 (0.43-2.34) | 0.53 (0.21-1.33) | |
| 11-20 times | 1.39 (0.65-2.97) | 0.41 (0.09-2.00) | 1.60 (0.68-3.75) | 0.38 (0.11-1.37) | |
| 21-50 times | 1.18 (0.42-3.26) | 0.78 (0.16-3.72) | 1.53 (0.53-4.49) | 0.73 (0.21-2.58) | |
| >50 times | 1.08 (0.24-4.78) | 1.76 (0.36-8.66) | 3.36 (1.16-9.71) | 1.13 (0.25-5.18) | 0.007 |
| **Sunburns as a youth (10-20 years old)** | | | | | |
| Never | Reference | Reference[3] | Reference | Reference | |
| 1-5 times | 2.21 (0.54-9.07) | - | 0.91 (0.29-2.86) | 1.44 (0.35-6.01) | |

| Variables | HR (95% CI)[2] Trunk (n=110) | Head/neck (n=48) | Upper limbs (n=83) | Lower limbs (n=64) | P-interaction |
|---|---|---|---|---|---|
| 6-10 times | 2.69 (0.64-11.27) | - | 1.20 (0.38-3.85) | 1.33 (0.31-5.69) | |
| 11-20 times | 3.58 (0.85-15.07) | - | 0.90 (0.26-3.12) | 0.57 (0.11-2.89) | |
| 21-50 times | 1.65 (0.32-8.59) | - | 1.80 (0.52-6.29) | 1.00 (0.18-5.49) | |
| >50 times | 3.32 (0.60-18.49) | - | 3.26 (0.87-12.22) | 2.02 (0.33-12.31) | <0.001 |
| **Sunburns as an adult (more than 20 years old)** | | | | | |
| Never | Reference | Reference | Reference | Reference | |
| 1-5 times | 2.59 (1.18-5.66) | 1.12 (0.47-2.66) | 1.12 (0.58-2.17) | 1.07 (0.53-2.20) | |
| 6-10 times | 3.42 (1.48-7.89) | 1.84 (0.71-4.81) | 0.66 (0.26-1.68) | 0.93 (0.38-2.26) | |
| 11-20 times | 2.17 (0.81-5.82) | 0.57 (0.07-4.83) | 1.51 (0.63-3.62) | 0.98 (0.33-2.87) | |
| 21-50 times | 3.26 (1.09-9.76) | 0.53 (0.06-4.51) | 3.68 (1.58-8.57) | 1.71 (0.51-5.69) | |
| >50 times | 3.97 (1.01-15.56) | - | 1.90 (0.42-8.58) | 1.18 (0.15-9.31) | <0.001 |
| **Cumulative sun exposure**[4] | | | | | |
| Q1 | Reference | Reference | Reference | Reference | |
| Q2 | 1.38 (0.71-2.70) | 1.14 (0.40-3.24) | 0.63 (0.34-1.15) | 0.68 (0.33-1.40) | |
| Q3 | 1.45 (0.76-2.78) | 1.81 (0.70-4.71) | 0.58 (0.31-1.09) | 0.53 (0.24-1.18) | |
| Q4 | 1.13 (0.59-2.17) | 1.38 (0.51-3.68) | 0.50 (0.26-0.93) | 1.26 (0.64-2.47) | 0.89 |
| **Average number of hours in the sun on weekdays**[4] | | | | | |
| Q1 | Reference | Reference | Reference | Reference | |
| Q2 | 1.49 (0.79-2.83) | 1.78 (0.62-5.12) | 0.70 (0.39-1.26) | 0.93 (0.47-1.86) | |
| Q3 | 1.46 (0.78-2.73) | 1.98 (0.73-5.41) | 0.51 (0.27-0.97) | 0.67 (0.32-1.39) | |
| Q4 | 1.23 (0.66-2.27) | 2.01 (0.74-5.41) | 0.53 (0.28-0.99) | 0.89 (0.43-1.83) | 0.43 |
| **Average number of hours in the sun on weekend days**[4] | | | | | |
| Q1 | Reference | Reference | Reference | Reference | |
| Q2 | 1.99 (1.10-3.62) | 0.89 (0.34-2.32) | 0.73 (0.39-1.37) | 0.92 (0.45-1.88) | |
| Q3 | 1.17 (0.62-2.22) | 1.02 (0.43-2.40) | 0.79 (0.43-1.44) | 1.01 (0.50-2.03) | |
| Q4 | 1.49 (0.80-2.79) | 1.30 (0.55-3.11) | 0.76 (0.39-1.49) | 1.05 (0.49-2.24) | 0.22 |
| **Sunbed use** | | | | | |
| Never | Reference | Reference | Reference | Reference | |
| Ever | 1.28 (0.63-2.57) | 1.30 (0.47-3.57) | 1.74 (0.90-3.37) | 1.15 (0.53-2.51) | 0.65 |
| **Number of skin cancers excised** | | | | | |
| None | Reference | Reference | Reference | Reference | |
| 1 | 1.17 (0.63-2.19) | 2.22 (1.01-4.85) | 2.89 (1.51-5.53) | 0.57 (0.23-1.45) | |
| 2-10 | 1.97 (1.26-3.07) | 1.86 (0.89-3.88) | 3.31 (1.87-5.86) | 1.19 (0.66-2.15) | |
| 10-20 | 2.42 (1.05-5.56) | 1.46 (0.30-7.13) | 5.60 (2.36-13.26) | 1.62 (0.49-5.39) | |
| 20+ | 3.40 (1.50-7.69) | 1.82 (0.41-8.02) | 0.95 (0.12-7.52) | 3.10 (1.06-9.08) | <0.001 |

| Variables | HR (95% CI)[2] Trunk (n=110) | Head/neck (n=48) | Upper limbs (n=83) | Lower limbs (n=64) | P-interaction |
|---|---|---|---|---|---|
| **Number of actinic lesions burnt/frozen off** | | | | | |
| None | Reference | Reference | Reference | Reference | |
| 1-5 | 1.44 (0.84-2.45) | 1.78 (0.78-4.09) | 3.21 (1.52-6.80) | 1.13 (0.57-2.21) | |
| 6-10 | 2.13 (1.15-5.41) | 1.74 (0.60-5.07) | 4.46 (1.90-10.47) | 1.06 (0.40-2.80) | |
| 11-20 | 1.56 (0.76-3.18) | 1.30 (0.39-4.37) | 6.49 (2.89-14.56) | 2.59 (1.25-5.36) | |
| 21-50 | 2.19 (1.13-4.24) | 2.24 (0.71-7.02) | 6.38 (2.73-14.89) | 3.52 (1.67-7.40) | |
| 50+ | 2.40 (1.15-3.98) | 2.44 (0.82-7.27) | 6.60 (2.70-16.13) | 1.77 (0.62-5.06) | <0.001 |

[1]11 melanoma cases had missing information on site.
[2]Born in Australia, latitude of birth, region of Australia lived longest (as a child; lifetime) adjusted for age and sex.
All other variables adjusted for age, sex, hair colour and tanning ability.
[3]Zero cells in referent category.
[4]Q - Quartile

**Table S5.** Measures of **s**un exposure and risk of invasive melanoma, stratified by sex.

| | HR (95% CI)[1] | | |
|---|---|---|---|
| **Variables** | **Women** (n=136) | **Men** (n=180) | **P-interaction** |
| **Born in Australia** | | | |
| No | Reference | Reference | |
| Yes | 2.06 (1.17-3.63) | 1.29 (0.86-1.93) | 0.17 |
| **Latitude of birth** | | | |
| ≥45º S/N | Reference | Reference | |
| <45º S/N | 1.89 (0.97-3.70) | 1.48 (0.90-2.44) | 0.54 |
| **Region of Australia lived longest up to age 20 years** | | | |
| Southern region | Reference | Reference | |
| Central region | 1.15 (0.78-1.69) | 1.12 (0.80-1.56) | |
| Northern region | 0.62 (0.33-1.15) | 0.88 (0.54-1.43) | 0.76 |
| **Region of Australia lived longest over lifetime** | | | |
| Southern region | Reference | Reference | |
| Central region | 1.07 (0.71-1.61) | 1.05 (0.74-1.50) | |
| Northern region | 0.63 (0.34-1.16) | 0.76 (0.47-1.25) | 0.79 |
| **Age moved to Australia** | | | |
| Born in Australia | Reference | Reference | |
| 1-10 years | 0.42 (0.13-1.33) | 0.95 (0.49-1.86) | |
| 11-20 years | 0.66 (0.21-2.08) | 1.60 (0.81-3.15) | |
| 20+ years | 0.47 (0.22-1.00) | 0.48 (0.26-0.92) | 0.37 |
| **Sunburns as a child (less than 10 years)** | | | |
| Never | Reference | Reference | |
| 1-5 times | 1.21 (0.77-1.913) | 0.93 (0.60-1.43) | |
| 6-10 times | 1.05 (0.59-1.88) | 0.84 (0.49-1.44) | |
| 11-20 times | 0.91 (0.43-1.92) | 0.95 (0.51-1.75) | |
| 21-50 times | 0.38 (0.09-1.63) | 1.35 (0.69-2.65) | |
| >50 times | 1.83 (0.70-4.82) | 1.78 (0.76-4.19) | 0.40 |
| **Sunburns as a youth (10-20 years old)** | | | |
| Never | Reference | Reference | |
| 1-5 times | 1.56 (0.52-4.71) | 1.64 (0.66-4.08) | |
| 6-10 times | 2.05 (0.67-6.28) | 1.58 (0.62-4.01) | |
| 11-20 times | 1.18 (0.36-3.84) | 1.96 (0.76-5.04) | |
| 21-50 times | 1.21 (0.33-4.46) | 1.71 (0.60-4.84) | |
| >50 times | 2.36 (0.59-9.46) | 3.74 (1.29-10.87) | 0.43 |
| **Sunburns as an adult (more than 20 years old)** | | | |
| Never | Reference | Reference | |
| 1-5 times | 1.41 (0.85-2.33) | 1.35 (0.81-2.23) | |
| 6-10 times | 1.17 (0.61-2.24) | 1.70 (0.97-2.979) | |
| 11-20 times | 1.08 (0.47-2.45) | 1.41 (0.74-2.70) | |
| 21-50 times | 2.00 (0.80-5.01) | 2.62 (1.33-5.13) | |
| >50 times | 2.09 (0.49-8.98) | 1.56 (0.52-4.62) | 0.67 |
| **Cumulative sun exposure**[2] | | | |
| Q1 | Reference | Reference | |
| Q2 | 0.76 (0.49-1.18) | 1.13 (0.62-2.04) | |
| Q3 | 0.78 (0.49-1.25) | 1.34 (0.77-2.33) | |

| Variables | HR (95% CI)[1] Women (n=136) | Men (n=180) | P-interaction |
|---|---|---|---|
| Q4 | 0.62 (0.32-1.18) | 1.20 (0.70-2.03) | 0.20 |
| **Average number of hours in the sun on weekdays**[2] | | | |
| Q1 | Reference | Reference | |
| Q2 | 0.86 (0.56-1.32) | 1.38 (0.79-2.42) | |
| Q3 | 0.74 (0.47-1.18) | 1.36 (0.80-2.29) | |
| Q4 | 0.55 (0.27-1.09) | 1.32 (0.80-2.17) | 0.12 |
| **Average number of hours in the sun on weekend days**[2] | | | |
| Q1 | Reference | Reference | |
| Q2 | 1.05 (0.68-1.61) | 1.21 (0.70-2.07) | |
| Q3 | 0.67 (0.39-1.14) | 1.28 (0.77-2.13) | |
| Q4 | 1.01 (0.57-1.80) | 1.22 (0.74-2.02) | 0.37 |
| **Sunbed use** | | | |
| Never | Reference | Reference | |
| Ever | 1.47 (0.93-2.34) | 1.02 (0.50-2.08) | 0.36 |
| **Number of skin cancers excised** | | | |
| None | Reference | Reference | |
| 1 | 1.67 (1.04-2.67) | 1.24 (0.75-2.06) | |
| 2-10 | 1.86 (1.23-2.82) | 2.16 (1.52-3.08) | |
| 10-20 | 3.35 (1.56-7.20) | 2.85 (1.56-5.20) | |
| 20+ | 0.74 (0.11-5.28) | 3.20 (1.71-6.00) | 0.49 |
| **Number of actinic lesions burnt/frozen off** | | | |
| None | Reference | Reference | |
| 1-5 | 1.97 (1.22-3.20) | 1.56 (1.01-2.41) | |
| 6-10 | 2.33 (1.25-4.34) | 1.96 (1.17-3.29) | |
| 11-20 | 3.35 (1.85-6.06) | 2.00 (1.18-3.28) | |
| 21-50 | 3.74 (1.97-7.10) | 2.76 (1.68-4.52) | |
| 50+ | 3.73 (1.85-7.50) | 2.86 (1.70-4.82) | 0.92 |

[1]Born in Australia, latitude of birth, region of Australia lived longest (as a child; lifetime) adjusted for age and sex. All other variables adjusted for age, sex, hair colour and tanning ability.
[2]Q - Quartile

**Table S6.** Measures of sun exposure and risk of first melanoma (invasive or *in situ*).

| | Melanoma (invasive or *in situ*) | | |
|---|---|---|---|
| | **No** (n=38,050) | **Yes** (n=782) | |
| **Variables** | **n (%)** | **n (%)** | **HR (95%CI)[1]** |
| **Born in Australia** | | | |
| No | 31,334 (82.4) | 675 (86.3) | Reference |
| Yes | 6716 (17.7) | 107 (11.7) | 1.43 (1.17-1.76) |
| **Latitude of birth** | | | |
| ≥45º S/N | 4462 (11.7) | 67 (8.6) | Reference |
| <45º S/N | 33,588 (88.3) | 715 (91.4) | 1.54 (1.20-1.98) |
| **Region of Australia lived longest up to age 20 years** | | | |
| Southern region | 11,502 (33.8) | 245 (33.8) | Reference |
| Central region | 16,813 (49.4) | 379 (52.3) | 1.10 (0.93-1.29) |
| Northern region | 5749 (16.9) | 101 (13.9) | 0.87 (0.69-1.10) |
| **Region of Australia lived longest over lifetime** | | | |
| Southern region | 8677 (22.8) | 173 (22.1) | Reference |
| Central region | 22,602 (59.4) | 499 (63.8) | 1.12 (0.95-1.34) |
| Northern region | 6771 (17.8) | 110 (14.1) | 0.83 (0.65-1.05) |
| **Age moved to Australia** | | | |
| Born in Australia | 31,374 (82.5) | 675 (86.3) | Reference |
| 1-10 years | 1856 (4.9) | 30 (3.8) | 0.74 (0.51-1.07) |
| 11-20 years | 1079 (2.8) | 25 (3.2) | 1.02 (0.68-1.52) |
| 20+ years | 3741 (9.8) | 52 (6.7) | 0.60 (0.45-0.79) |
| **Sunburns as a child (less than 10 years)** | | | |
| Never | 8006 (21.0) | 133 (17.0) | Reference |
| 1-5 times | 17,380 (45.7) | 351 (44.9) | 1.08 (0.88-1.33) |
| 6-10 times | 6635 (17.4) | 151 (19.3) | 1.18 (0.92-1.51) |
| 11-20 times | 3585 (9.4) | 80 (10.2) | 1.22 (0.91-1.63) |
| 21-50 times | 1720 (4.5) | 43 (5.5) | 1.25 (0.88-1.78) |
| >50 times | 724 (1.9) | 24 (3.1) | 1.64 (1.05-2.57) |
| **Sunburns as a youth (10-20 years old)** | | | |
| Never | 1775 (2.7) | 22 (2.8) | Reference |
| 1-5 times | 16,615 (43.7) | 311 (39.8) | 1.56 (1.01-2.42) |
| 6-10 times | 9446 (24.8) | 216 (27.6) | 1.96 (1.26-3.04) |
| 11-20 times | 6329 (16.6) | 140 (17.9) | 1.85 (1.17-2.91) |
| 21-50 times | 2849 (7.5) | 58 (7.4) | 1.67 (1.01-2.74) |
| >50 times | 1036 (2.7) | 35 (4.5) | 2.68 (1.56-4.59) |
| **Sunburns as an adult (more than 20 years old)** | | | |
| Never | 6160 (16.2) | 111 (14.2) | Reference |
| 1-5 times | 20,236 (53.2) | 389 (49.7) | 1.21 (0.97-1.52) |
| 6-10 times | 6293 (16.5) | 160 (20.5) | 1.56 (1.21-2.00) |
| 11-20 times | 3374 (8.9) | 63 (8.1) | 1.12 (0.81-1.54) |
| 21-50 times | 1438 (3.8) | 46 (5.9) | 1.81 (1.27-2.58) |
| >50 times | 551 (1.5) | 13 (1.7) | 1.32 (0.74-2.35) |
| **Cumulative sun exposure[2]** | | | |

| | Melanoma (invasive or *in situ*) | | |
|---|---|---|---|
| | **No** (n=38,050) | **Yes** (n=782) | |
| **Variables** | **n (%)** | **n (%)** | **HR (95%CI)**[1] |
| Q1 | 9228 (24.3) | 185 (23.7) | Reference |
| Q2 | 9569 (25.2) | 172 (22.0) | 0.82 (0.66-1.02) |
| Q3 | 9617 (25.3) | 196 (25.1) | 0.82 (0.66-1.01) |
| Q4 | 9636 (25.3) | 229 (29.3) | 0.81 (0.65-1.00) |
| **Average number of hours in the sun on weekdays**[2] | | | |
| Q1 | 8920 (23.4) | 184 (23.5) | Reference |
| Q2 | 9333 (24.5) | 186 (23.8) | 0.96 (0.77-1.19) |
| Q3 | 10,032 (26.4) | 204 (26.1) | 0.89 (0.72-1.10) |
| Q4 | 9765 (25.7) | 208 (26.6) | 0.83 (0.67-1.02) |
| **Average number of hours in the sun on weekend days**[2] | | | |
| Q1 | 9589 (25.2) | 182 (23.3) | Reference |
| Q2 | 9724 (24.2) | 217 (27.8) | 1.17 (0.95-1.44) |
| Q3 | 9652 (25.4) | 197 (25.2) | 1.00 (0.81-1.24) |
| Q4 | 9085 (23.9) | 186 (23.8) | 0.97 (0.78-1.22) |
| **Sunbed use** | | | |
| Never | 33,976 (89.3) | 710 (90.8) | Reference |
| Ever | 4074 (10.7) | 72 (9.2) | 1.18 (0.92-1.51) |
| **Number of skin cancers excised** | | | |
| None | 23,366 (61.4) | 292 (37.3) | Reference |
| 1 | 5291 (13.9) | 122 (15.6) | 1.71 (1.38-2.11) |
| 2-10 | 7875 (20.7) | 286 (36.6) | 2.38 (2.00-2.82) |
| 10-20 | 917 (2.4) | 46 (5.9) | 2.84 (2.07-3.91) |
| 20+ | 601 (1.6) | 36 (4.6) | 3.11 (2.16-4.47) |
| **Number of actinic lesions burnt/frozen off** | | | |
| None | 17,187 (45.2) | 192 (24.6) | Reference |
| 1-5 | 10,317 (27.1) | 192 (24.6) | 1.55 (1.27-1.90) |
| 6-10 | 3595 (9.5) | 114 (14.6) | 2.37 (1.87-3.019) |
| 11-20 | 2904 (9.5) | 98 (12.5) | 2.39 (1.87-3.07) |
| 21-50 | 2215 (5.8) | 89 (11.4) | 2.69 (2.08-3.49) |
| 50+ | 1832 (4.8) | 97 (12.4) | 3.23 (2.48-4.21) |

[1]Born in Australia, latitude of birth, region of Australia lived longest (as a child; lifetime) adjusted for age.
All other variables adjusted for age, hair colour and tanning ability.
[2]Q - Quartile
